# Association between body mass index (BMI) and hypertension in South Asian population: Evidence from Demographic and Health Survey

**DOI:** 10.1101/605469

**Authors:** Fariha Binte Hossain, Shajedur Rahman Shawon, Gourab Adhikary, Arif Chowdhury

**Author notes:** **CORRESPONDING AUTHOR:** Md Shajedur Rahman Shawon, Cancer Epidemiology Unit, Nuffield Department of Population Health, University of Oxford, Richard Doll Building, OX3 7LF, Oxford, UK. These authors contributed equally to this work (Fariha Binte Hossain, Shajedur Rahman Shawon).

## Abstract

Although there has been a well-established association between adiposity and hypertension, whether such associations are heterogeneous for South Asian populations or for different socioeconomic groups is not well-known. We analysed the recent Demographic and Health Survey (DHS) data from Bangladesh, India, and Nepal to estimate the age-specific prevalence of hypertension and the association of body mass index (BMI) with hypertension. We used multiple logistic regressions to estimate the odds ratios (ORs) with 95% confidence intervals (CIs) of hypertension for overweight and obesity as well as for each 5-unit increase in BMI. The overall prevalence for hypertension among participants aged 35-44 years were 17.4%, 20%, and 22.5% for Bangladesh, India, and Nepal, respectively. For all age groups, men were more likely to be hypertensive than women in India and Nepal, but not in Bangladesh. Overweight and obesity were associated with higher odds of hypertension in all countries. For each 5 kg/m^2^ increase in BMI, the ORs for hypertension were 1.79 (95% CI: 1.65-1.93), 1.59 (95% CI: 1.58-1.61), and 2.03 (95% CI: 1.90-2.16) in Bangladesh, India, and Nepal, respectively. The associations between BMI and hypertension were consistent across various subgroups defined by sex, age, urbanicity, educational attainment and household’s wealth index. Our study shows that the association of BMI with hypertension is stronger for South Asian populations, and public health measures to reduce population-level reduction in BMI would also help in lowering the burden of hypertension.

## INTRODUCTION

Hypertension is one of the important preventable noncommunicable disease (NCD) risk factors for premature death and disability.^1–3^ About one-third of world’s adult population are hypertensive – according to recent reports.^4,5^ The burden of hypertension is increasing particularly in the low- and middle-income countries (LMICs).^5^ South Asia comprises of several LMICs and almost one-quarter of the world’s population lives in South Asia. Therefore, a greater understanding of the burden of hypertension in this region is required to develop public health interventions to control it.

There has been a well-established association between adiposity and hypertension in developed settings,^6–8^ but whether such association is heterogeneous for South Asian population is not well-known. Assessing association between body mass index (BMI) and hypertension has important public health implications in South Asian countries, where the burden of hypertension is high and obesity is increasing at the population level.^9–11^ In addition, looking at the association in subgroups defined by sex, age, urbanicity, and socioeconomic status is crucial to understand how consistent the association between BMI and hypertension is across different groups. There is no study, to the best of our knowledge, which looked at the association of BMI with hypertension across different groups in nationally-representative samples from South Asian countries.

In this study, we aim to examine the age-specific prevalence of hypertension in three South Asian countries, namely Bangladesh, India, and Nepal. We also systematically assess the association between overweight-obesity and hypertension using different cut-offs, and how the association between BMI and hypertension varies across a wide variety of subgroups of population.

## METHODS

### Study design and data sources

This study is based on three South Asian countries, namely Bangladesh, India, and Nepal. Recent Demographic and Health Survey (DHS) data for these countries had information on both blood pressure and anthropometry for adult population.

DHS are periodical nationally-representative household surveys which provide data for a wide range of variables on population, health, and nutrition. These surveys usually are conducted by a national implementing agency with technical assistances provided by the DHS program. Surveys are based on two-stage stratified sampling of households – firstly, sampling census enumeration areas are selected using probability proportional to size (PPS) sampling technique through statistics provided by the respective national statistical office, and secondly, households are selected through systematic random sampling from the complete listing of households within a selected enumeration area. From these selected households, subsamples of eligible participants are additionally selected for biomarker testing, which includes height, weight, and blood pressure.^12^

DHS surveys receive ethical approval both from the ICF Institutional Review Board and from a country-specific review board. Informed consent is taken from each participant for their participation in the survey and for anthropometric and blood pressure measurements. The DHS program authorises researchers to use relevant datasets for analysis upon submission of a brief research proposal. The data we received for this study were anonymized for protection of privacy, anonymity and confidentiality. More details on survey design, ethical approval, data availability can be found in the DHS program website [https://dhsprogram.com/].

We included those who had consented for measurement of blood pressure, height, and weight, as well as had valid information for those variables. DHS surveys have very high response rate, usually more than 90%. We used the household member record dataset which has one record for every household member, and includes variables for sociodemographic, height, weight and blood pressure measurement.

### Anthropometric measurement and BMI classification

In the included DHS surveys, height and weight of the participants were measured by trained personnel using standardized instruments and procedures. BMI was then calculated by dividing body weight (kg) by squared height (m^2^). We classified participants based into four groups according to the conventional World Health Organization (WHO) classification system:^13^ underweight (<18.5 kg/m^2^), normal weight (18.5-24.9 kg/m^2^), overweight (25.0-29.9 kg/m^2^), and obese (≥30.0 kg/m^2^). We also classified them according to the proposed cut-offs for South Asian population: underweight (<18.0 kg/m^2^), normal weight (18.0-22.9 kg/m^2^), overweight (23.0-27.4 kg/m^2^) and obese (≥27.5 kg/m^2^).^14^

### Blood pressure measurement and hypertension

Blood pressure was measured for participants using a standard protocol.^15^ In brief, three measurements were taken by trained health technicians, at seating position, at about 10 minutes intervals. The mean of the second and third measurement was used to record systolic blood pressure and diastolic blood pressure.

We defined hypertension based on the cut-offs provided by the Seventh Report of Joint National Committee on Prevention, Detection, Evaluation, and Treatment of High Blood Pressure (JNC7) guideline 2003^16^ and also the 2017 American College of Cardiology/American Heart Association (2017 ACC/AHA) Guideline for the Prevention, Detection, Evaluation, and Management of High Blood Pressure in Adults.^17^ According to the JNC7, an individual was categorised as hypertensive if they had systolic blood pressure ≥140 mmHg or diastolic blood pressure ≥90 mmHg or reported about antihypertensive medication use during the survey. According to the 2017 ACC/AHA, an individual was categorised as hypertensive if they had systolic blood pressure ≥130 mmHg or diastolic blood pressure ≥80 mmHg or reported about antihypertensive medication use during the survey.

### Other covariates

DHS collected information on wide range variables from the selected households and the respondents from those households using face-to-face interview conducted by trained personnel. DHS collected information on socioeconomic factors like area of residence and household’s wealth index. Place of residence (rural and urban) was defined according to country-specific definitions. For household’s wealth index, each national implementing agency constructed a country-specific index using principal components analysis from data on household assets including durable goods (i.e. bicycles, televisions etc.) and dwelling characteristics (i.e. sanitation, source of drinking water and construction material of house etc.).^12^ This wealth index was then categorized into five groups (i.e. poorest, poorer, middle, richer, and richest) based on the quintile distribution of the sample.

### Statistical analyses

All analyses were conducted following the instructions in the DHS guide to analysis.^18^ All analyses were performed using Stata v15.1 (Statacorp, College Station, TX, USA). Considering the two-stage stratified cluster sampling in DHS, we applied Stata’s survey estimation procedures (“*svy*” command) for estiamations.^19^ We looked at the descriptive statistics by sex on sociodemographic, anthropometric, and blood pressure variables using proportions for categorical variables and mean and standard deviation (SD) for continuous variables. We used sampling weights given in each DHS dataset in order to get nationally-representative estimates. 95% confidence intervals (CIs) for prevalence estimates were calculated using a logit transform of the estimate.

To examine the association between BMI and hypertension, we used multiple logistic regressions, separately for each included country. We also estimated the trend by estimating the odds ratios (ORs) with 95% confidence intervals (CIs) of hypertension for each 5 kg/m^2^ increase in BMI. All these analyses were adjusted for age, sex, are of residence, household’s highest education level, and household’s wealth index, as appropriate. We then examined the trend in subgroups of individuals defined by various characteristics.

## RESULTS

A total of 821 040 men and women from Bangladesh, India, and Nepal were included in this study. Table 1 shows that sociodemographic characteristics for three study population, by sex. Study populations varied widely for age – mean age for participants from Bangladesh was 51 years whereas the mean ages for participants from other two countries were much lower (India: 30 years and Nepal: 38 years). Almost two-thirds of the participants were from rural areas in Bangladesh and India, but Nepal had more participants from urban areas. Male participants were more likely to be educated than female participants in all countries, and India had higher proportions of men and women educated to secondary or higher level. Wealth index distributions were similar between men and women, and also among countries (Table 1).

**Table 1:** Sociodemographic characteristics of three study populations, by sex

|  | Bangladesh |  | India |  | Nepal |  |
| --- | --- | --- | --- | --- | --- | --- |
|  | Male | Female | Male | Female | Male | Female |
| Number of participants | 3798 | 3837 | 109527 | 689131 | 6114 | 8633 |
| Age in years, mean (SD) | 51.7 (12.9) | 50.4 (12.5) | 31.7 (11.1) | 29.8 (9.8) | 40.1 (18.2) | 36.9 (16.9) |
| Type of place of residence, n (%) |  |  |  |  |  |  |
| Urban | 1253 (33.0) | 1254 (32.7) | 34137 (31.2) | 199227 (28.9) | 3884 (63.5) | 5459 (63.2) |
| Rural | 2545 (67.0) | 2583 (67.3) | 75390 (68.8) | 489904 (71.1) | 2230 (36.5) | 3174 (36.8) |
| Highest educational level attained, n (%) |  |  |  |  |  |  |
| No education, preschool | 1330 (35.0) | 2112 (55.0) | 13874 (12.7) | 186695 (27.1) | 1428 (23.4) | 4089 (47.4) |
| Primary | 1076 (28.3) | 1035 (27.0) | 14250 (13.0) | 93170 (13.5) | 1278 (20.9) | 1141 (13.2) |
| Secondary | 890 (23.4) | 538 (14.0) | 64081 (58.5) | 330514 (48.0) | 2398 (39.2) | 2440 (28.3) |
| Higher | 502 (13.2) | 152 (4.0) | 17064 (15.6) | 77464 (11.2) | 1005 (16.4) | 959 (11.1) |
| Wealth index, n (%) |  |  |  |  |  |  |
| Poorest | 681 (17.9) | 664 (17.3) | 18302 (16.7) | 132389 (19.2) | 1293 (21.1) | 1917 (22.2) |
| Poorer | 702 (18.5) | 686 (17.9) | 22874 (20.9) | 147995 (21.5) | 1241 (20.3) | 1792 (20.8) |
| Middle | 732 (19.3) | 750 (19.5) | 23782 (21.7) | 145007 (21.0) | 1185 (19.4) | 1754 (20.3) |
| Richer | 769 (20.2) | 816 (21.3) | 22620 (20.7) | 135960 (19.7) | 1276 (20.9) | 1738 (20.1) |
| Richest | 914 (24.1) | 921 (24.0) | 21949 (20.0) | 127780 (18.5) | 1119 (18.3) | 1432 (16.6) |

Table 2 shows the distribution of anthropometric and blood pressure measurements for the study populations. On average, females had slightly higher BMI than males. According to both the WHO classification and South Asian classification systems, more women were overweight and obese in all three countries. In Bangladesh, women had higher systolic (mean 121.0 vs 116.2 mmHg) and diastolic (mean 79.6 vs 76.4 mmHg) blood pressure than men. In contrary, men had higher mean blood pressure than women in India (systolic: 121.8 vs 115.2 mmHg; diastolic: 79.9 vs 78.1 mmHg) and Nepal (systolic: 120.0 vs. 112.4 mmHg; diastolic 79.0 vs. 76.4 mmHg) (Table 2).

**Table 2:**
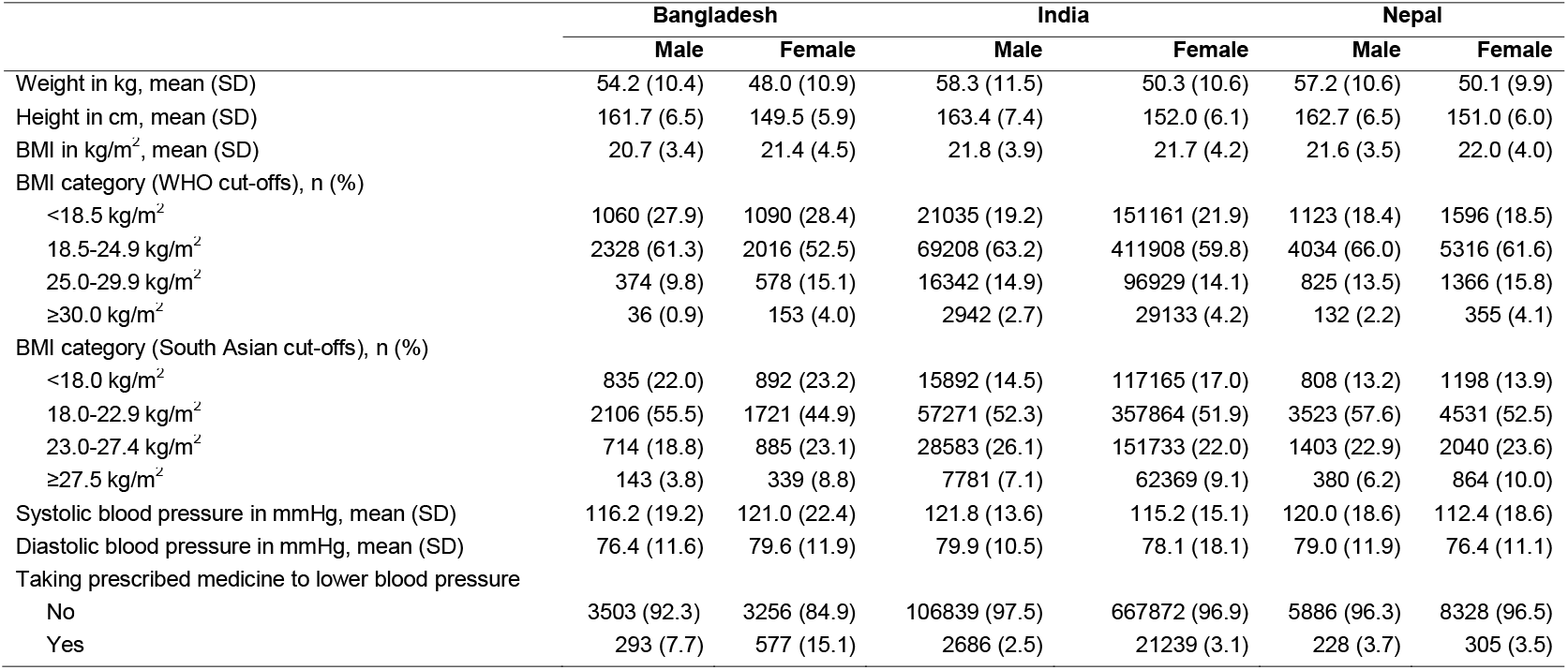
Distribution of anthropometric and blood pressure measurements among the three study populations, by sex

|  | Bangladesh |  | India |  | Nepal |  |
| --- | --- | --- | --- | --- | --- | --- |
|  | Male | Female | Male | Female | Male | Female |
| Weight in kg, mean (SD) | 54.2 (10.4) | 48.0 (10.9) | 58.3 (11.5) | 50.3 (10.6) | 57.2 (10.6) | 50.1 (9.9) |
| Height in cm, mean (SD) | 161.7 (6.5) | 149.5 (5.9) | 163.4 (7.4) | 152.0 (6.1) | 162.7 (6.5) | 151.0 (6.0) |
| BMI in kg/m<sup>2</sup>, mean (SD) | 20.7 (3.4) | 21.4 (4.5) | 21.8 (3.9) | 21.7 (4.2) | 21.6 (3.5) | 22.0 (4.0) |
| BMI category (WHO cut-offs), n (%) |  |  |  |  |  |  |
| <18.5 kg/m<sup>2</sup> | 1060 (27.9) | 1090 (28.4) | 21035 (19.2) | 151161 (21.9) | 1123 (18.4) | 1596 (18.5) |
| 18.5-24.9 kg/m<sup>2</sup> | 2328 (61.3) | 2016 (52.5) | 69208 (63.2) | 411908 (59.8) | 4034 (66.0) | 5316 (61.6) |
| 25.0-29.9 kg/m<sup>2</sup> | 374 (9.8) | 578 (15.1) | 16342 (14.9) | 96929 (14.1) | 825 (13.5) | 1366 (15.8) |
| ≥30.0 kg/m<sup>2</sup> | 36 (0.9) | 153 (4.0) | 2942 (2.7) | 29133 (4.2) | 132 (2.2) | 355 (4.1) |
| BMI category (South Asian cut-offs), n (%) |  |  |  |  |  |  |
| <18.0 kg/m<sup>2</sup> | 835 (22.0) | 892 (23.2) | 15892 (14.5) | 117165 (17.0) | 808 (13.2) | 1198 (13.9) |
| 18.0-22.9 kg/m<sup>2</sup> | 2106 (55.5) | 1721 (44.9) | 57271 (52.3) | 357864 (51.9) | 3523 (57.6) | 4531 (52.5) |
| 23.0-27.4 kg/m<sup>2</sup> | 714 (18.8) | 885 (23.1) | 28583 (26.1) | 151733 (22.0) | 1403 (22.9) | 2040 (23.6) |
| ≥27.5 kg/m<sup>2</sup> | 143 (3.8) | 339 (8.8) | 7781 (7.1) | 62369 (9.1) | 380 (6.2) | 864 (10.0) |
| Systolic blood pressure in mmHg, mean (SD) | 116.2 (19.2) | 121.0 (22.4) | 121.8 (13.6) | 115.2 (15.1) | 120.0 (18.6) | 112.4 (18.6) |
| Diastolic blood pressure in mmHg, mean (SD) | 76.4 (11.6) | 79.6 (11.9) | 79.9 (10.5) | 78.1 (18.1) | 79.0 (11.9) | 76.4 (11.1) |
| Taking prescribed medicine to lower blood pressure |  |  |  |  |  |  |
| No | 3503 (92.3) | 3256 (84.9) | 106839 (97.5) | 667872 (96.9) | 5886 (96.3) | 8328 (96.5) |
| Yes | 293 (7.7) | 577 (15.1) | 2686 (2.5) | 21239 (3.1) | 228 (3.7) | 305 (3.5) |

Overall, Bangladesh had higher prevalence of hypertension (both overall and by sex) than India and Nepal, but it is important to remember that Bangladesh had older study participants than the other two. When we looked at the age-specific prevalence of hypertension, there was a sharp increase in prevalence of hypertension by age (Figure 1). The overall prevalence for hypertension among participants aged 35-44 years were 17.4%, 20%, and 22.5% for Bangladesh, India, and Nepal, respectively. For age groups 45-54 years, the prevalence increased to 25% in Bangladesh, 28.6% in India, and 30% in Nepal. For all age groups, men had higher prevalence of hypertension than women in India and Nepal, but not in Bangladesh. When we used the 2017 ACC/AHA guidelines to define hypertension, the prevalence estimates, as expected, increased significantly for all age groups in all three countries (Supplementary Figure S1).

**Figure 1:**
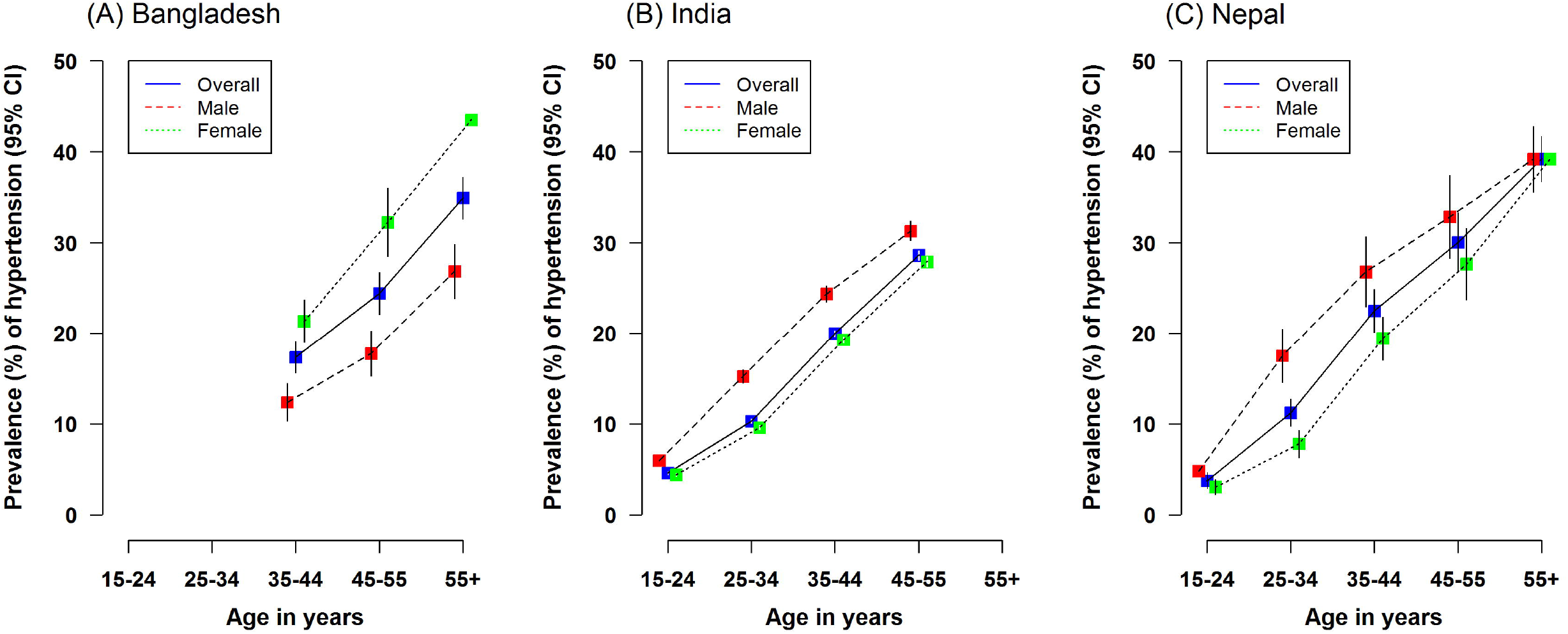
Age-specific prevalence of hypertension in three study populations, overall and by sex

After adjustment for five sociodemographic factors including age, sex, area of residence, wealth index, and highest educational attainment, being overweight and obese individuals, independent of classification system, had higher odds of having hypertension when compared to normal weight individuals (Table 3). Overweight people had almost two-fold increase in the odds of hypertension, whereas obese people had more than three-fold higher odds of hypertension. For each 5 kg/m^2^ increase in BMI, the ORs for hypertension were 1.79 (95% CI: 1.65-1.93), 1.59 (95% CI: 1.58-1.61), and 2.03 (95% CI: 1.90-2.16) in Bangladesh, India, and Nepal, respectively. We found similar associations between BMI and hypertension for all three countries when we used the AHA 2017 guidelines for defining hypertension (Supplementary Table S1).

**Table 3:** Adjusted odds ratios (ORs) with 95% CI for hypertension by BMI

|  | Bangladesh |  | India |  | Nepal |  |
| --- | --- | --- | --- | --- | --- | --- |
|  | No. of cases | OR (95% CI)<sup>†</sup> | No. of cases | OR (95% CI)<sup>†</sup> | No. of cases | OR (95% CI)<sup>†</sup> |
| **BMI categories (WHO cut-offs)** |  |  |  |  |  |  |
| Underweight (<18.5 kg/m<sup>2</sup>) | 383 | 0.57 (0.50-0.65) | 12121 | 0.70 (0.69-0.72) | 301 | 0.55 (0.48-0.63) |
| Normal weight (18.5-25.0 kg/m<sup>2</sup>) | 1110 | 1.00 (0.93-1.07) | 55386 | 1.00 (0.99-1.01) | 1519 | 1.00 (0.94-1.06) |
| Overweight (25.0-29.9 kg/m<sup>2</sup>) | 394 | 1.80 (1.57-2.07) | 27738 | 1.99 (1.96-2.02) | 757 | 2.46 (2.24-2.71) |
| Obese (≥30.0 kg/m<sup>2</sup>) | 108 | 2.72 (2.00-3.68) | 10762 | 3.03 (2.96-3.11) | 213 | 3.62 (2.97-4.41) |
| **BMI categories (South Asian cut-offs)** |  |  |  |  |  |  |
| Underweight (<18.0 kg/m<sup>2</sup>) | 322 | 0.73 (0.63-0.83) | 9221 | 0.79 (0.77-0.80) | 233 | 0.65 (0.56-0.76) |
| Normal weight (18.0-23.0 kg/m<sup>2</sup>) | 820 | 1.00 (0.92-1.09) | 40596 | 1.00 (0.99-1.01) | 1145 | 1.00 (0.93-1.07) |
| Overweight (23.0-27.0 kg/m<sup>2</sup>) | 614 | 2.14 (1.93-2.38) | 34799 | 1.76 (1.74-1.79) | 911 | 2.03 (1.87-2.20) |
| Obese (≥27.0 kg/m<sup>2</sup>) | 239 | 2.99 (2.46-3.64) | 21391 | 3.04 (2.99-3.10) | 501 | 3.64 (3.19-4.16) |
| **Trend (per 5 kg/m<sup>2</sup>)** | 1995 | 1.79 (1.65-1.93) | 106007 | 1.59 (1.58-1.61) | 2790 | 2.03 (1.90-2.16) |

To assess any further potential for effect modification by other factors, the OR per 5 kg/m^2^ was compared across subgroups of various individual characteristics, including sex, area of residence, age group, highest educational attainment, and household’s wealth index (Figure 2). Weak evidence of heterogeneity in the association between BMI and hypertension was found by sex (higher magnitude in males than females) in India and Nepal. For other characteristics, no significant heterogeneity was observed by subgroups consistently in three study populations.

**Figure 2:**
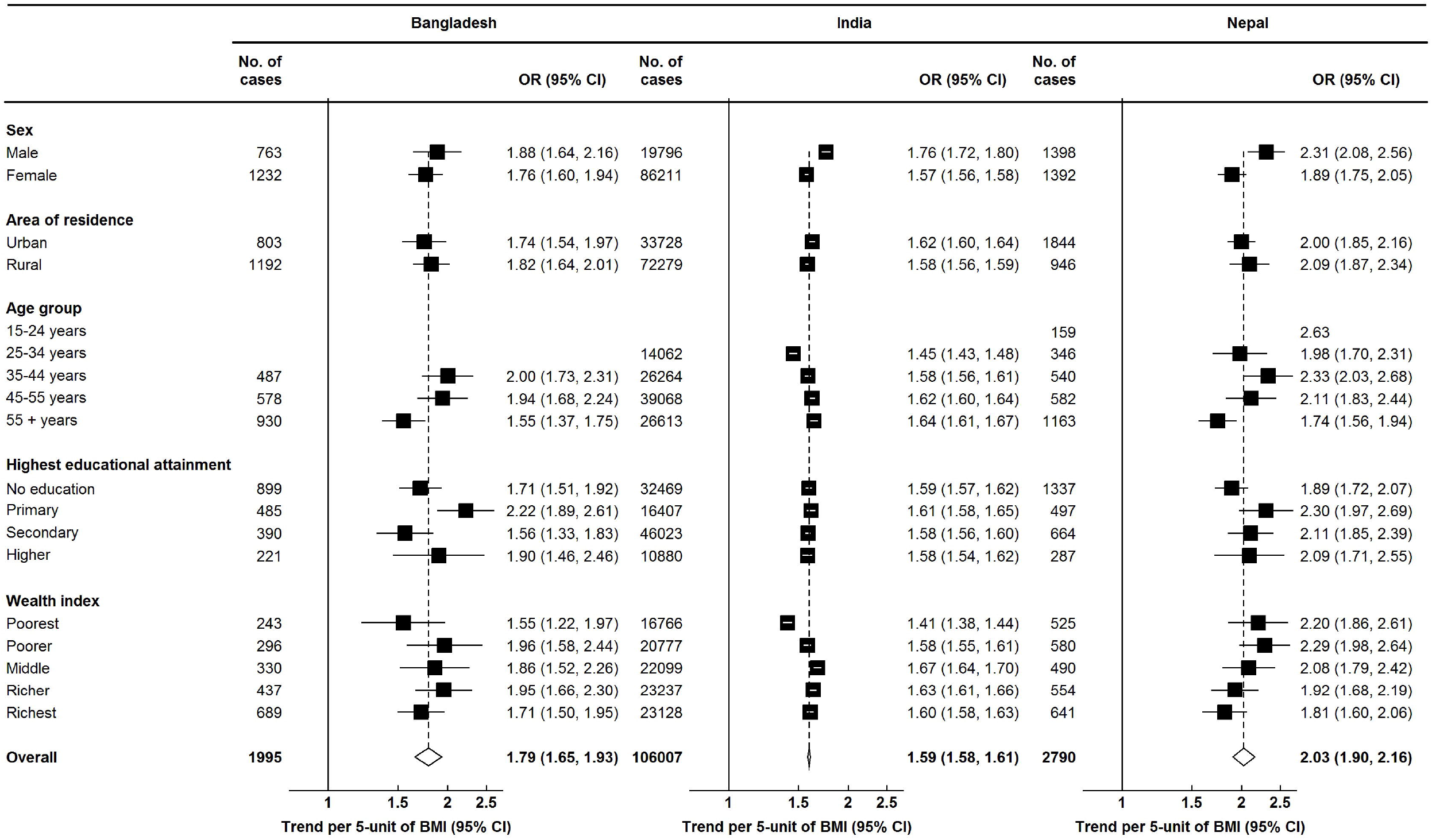
Odds ratios (ORs) with 95% confidence intervals (CIs) of hypertension per 5 kg/m^2^ increase in body mass index (BMI), by various characteristics Logistic regression models were adjusted for age, sex, area of residence, wealth index and highest educational attainment, as appropriate.

## DISCUSSION

This study involving more than 800 000 men and women from recent nationally-representative cross-sectional studies in three South Asian countries showed high prevalence of hypertension, particularly with increasing age. There were significant associations between overweight-obesity and hypertension, irrespective of cut-offs for defining overweight-obesity as well as hypertension. The associations between BMI and hypertension were consistent across various subgroups defined by sex, age, urbanicity, educational attainment and household’s wealth index, implying the robustness of such association.

Our study showed that almost one in every five adults aged 35 years and above in Bangladesh, India, and Nepal had hypertension. A recent systematic review^11^ showed considerable differences in prevalence estimates of hypertension in South Asian countries, but it did not consider the effects of differential age structure in the included studies. When we looked at the age-specific prevalence of hypertension, the country-specific prevalence estimates were almost similar. We found higher prevalence of hypertension among men than among women in India and Nepal, but not in Bangladesh. However, previous studies from this region mostly found the prevalence of hypertension was higher among men than among women.^20–24^ Systematic analysis of population-based studies from 90 countries showed that the age-standardized prevalence of hypertension between 2000 and 2010 decreased by 2.6% in developed countries, while in LMICs it increased by 7.7% during the same period.^5^ The high prevalence of hypertension in these three counties could be due to adoption of unhealthy lifestyles including intake of energy-dense foods, sedentary lifestyles, and rising level of obesity in the population.^5,10,20,24^ Positive associations between BMI and hypertension have been well reported in studies conducted among different ethnic groups.^6–8,25–28^ However, previous studies found that Asian populations had a much stronger association between BMI and blood pressure.^26,27^ Our study adds to the evidence suggesting that there are ethnic differences in the strength of the association between BMI and hypertension. South Asian populations may be at greater risk of developing hypertension with increasing BMI than any other ethnic groups.^28^ BMI has been found to be associated with hypertension, diabetes, and other NCDs in South Asian populations, at a much lower threshold level than the level for other populations.^20,21,29^ The possible reasons for such differences could be genetic and metabolic variations, as well as clustering of environmental, dietary, and social factors associated with hypertension.^9,27–29^ Previous studies looking at the relationship between adiposity and hypertension in this population were heterogeneous in terms of definitions used for overweight and obesity.^11,21–23^ We looked at the association using both WHO and South Asian cutoffs and also for each 5 kg/m^2^ increase in BMI. Our findings on the association between BMI and hypertension are consistent with previous literature.^6–8^

Additionally, we were able to show that this association is consistent across a wide range of subgroups defined by various characteristics. This means that the association between BMI and hypertension is more likely to be biological rather than environmental.

Since BMI is log-linearly associated with hypertension, any amount of reduction in BMI at population level can reduce the burden of hypertension at a large scale. Early diagnosis and treatment of hypertension is crucial for reducing NDC burden in South Asian countries,^23^ but given the robust and linear association between BMI and hypertension, primary prevention through reducing BMI would have much greater effect on reduction of cardiovascular morbidity and mortality. Previous studies found that awareness about high blood pressure and use of antihypertensive medication is low in this region.^22,23,30^ Also, the health systems are not well-prepared to manage the large burden of NDCs.^31,32^ Therefore, the policy makers should focus mainly on reduction of BMI at population level as one of the most important primary prevention strategies.

This study was limited by the use of cross-sectional data. There are possibilities of reverse causation; and we cannot establish a causal association between BMI and hypertension, or whether BMI is an independent risk factor of hypertension. We did not have dietary and lifestyle variables which could be potential mediators or could confound the observed associations. However, to the best of our knowledge, our study is the first to look at the association between BMI and hypertension in various subgroups of population. Taking advantage of the large sample size of our study, we were able to show that the associations of BMI with hypertension were robust across various socioeconomic subgroups. We also did several additional analyses using different cut-offs for defining both overweight-obesity and hypertension.

In conclusion, the age-specific prevalence of hypertension is very high among men and women in Bangladesh, India, and Nepal. The associations of BMI with hypertension are positive and robust across various subgroups of population defined by socioeconomic groups. Public health interventions targeting to reduce BMI at population level would have larger effects on reducing the burden of hypertension in South Asia.

## ACKNOWLEDGEMENTS

The authors thank the participants of Demographic and Health Surveys used in this study from Bangladesh, India, and Nepal. We would also like to thank the DHS Program to authorize us to use the data.

## AUTHOR CONTRIBUTIONS

Conception and design: FH, MS, GA, and AC

Data collection and management: FH, MS, and GA

Data analysis: FH, MS, GA

Interpretation of the results: All authors

Drafting of the article: FH and MS

Critical revision of the article for important intellectual content: All authors

Final approval of the article: All authors

## SUMMARY TABLE

### What is known?

- In many South Asian countries, there have been increasing trends of obesity and hypertension.
- Body mass index is positively associated with hypertension, but whether such association is consistent across socioeconomic subgroups is not clear.

### What this study adds?

- Almost one in five men and women aged 35 years and above in Bangladesh, India, and Nepal had hypertension.
- There were significant associations between overweight-obesity and hypertension, irrespective of cut-offs for defining overweight-obesity as well as hypertension.
- The associations between BMI and hypertension were consistent across various subgroups defined by sex, age, urbanicity, educational attainment and household’s wealth index.

