## Supplementary materials for "Association between body mass index (BMI) and hypertension in South Asian population: Evidence from Demographic and Health Survey"

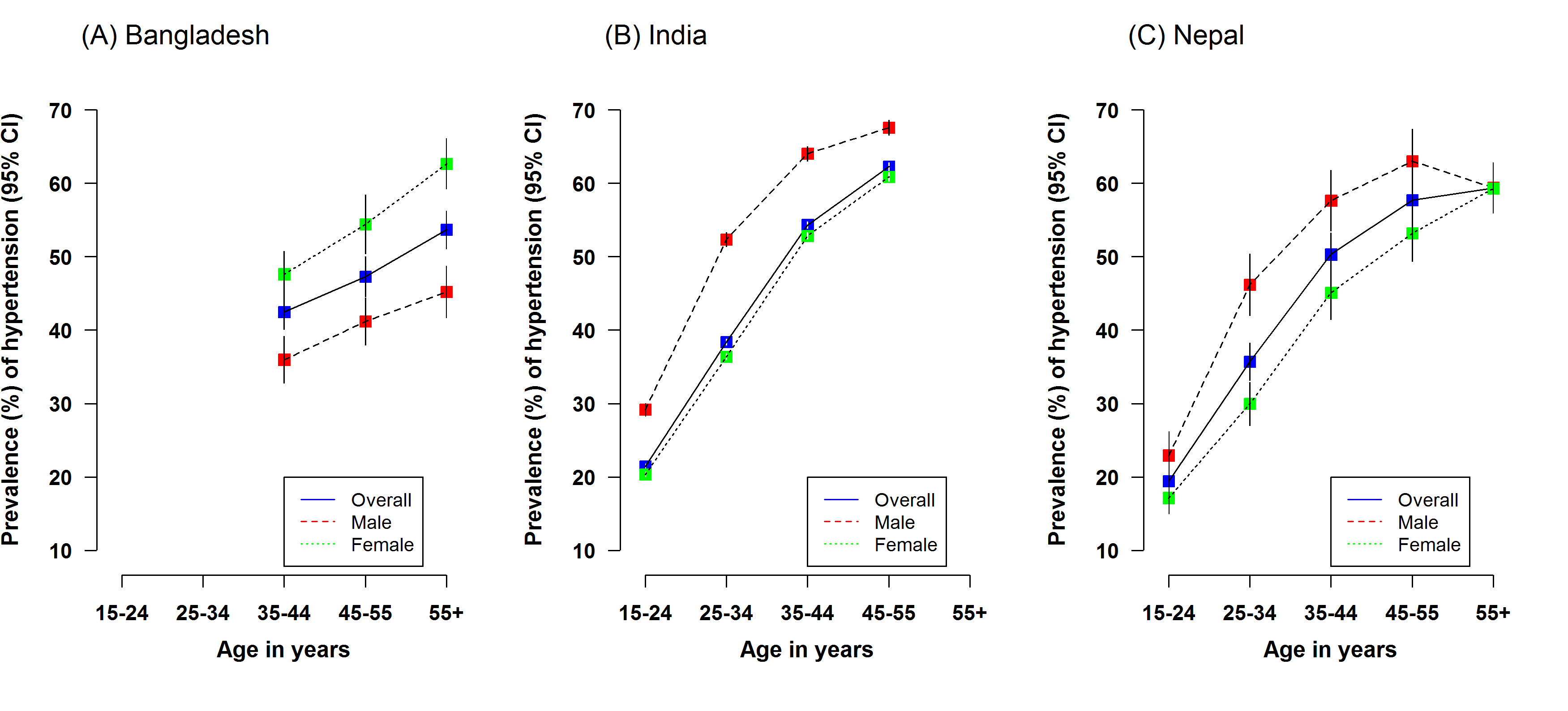

**Figure S1**: Prevalence of hypertension according to American Heart Association guideline, overall and by sex

**Table S1:** Adjusted odds ratios (ORs) with 95% CI for hypertension defined by the American Heart Association 2017 guideline, by BMI

|  | **Bangladesh** | | **India** | | **Nepal** | |
| --- | --- | --- | --- | --- | --- | --- |
|  | **No. of cases** | **OR (95% CI)** ^†^ | **No. of cases** | **OR (95% CI)** ^†^ | **No. of cases** | **OR (95% CI)** ^†^ |
| **BMI categories (WHO cut-offs)** |  |  |  |  |  |  |
| Underweight (<18.5 kg/m^2^) | 751 | 0.54 (0.49-0.60) | 44268 | 0.62 (0.61-0.63) | 725 | 0.52 (0.48-0.58) |
| Normal weight (18.5-25.0 kg/m^2^) | 2132 | 1.00 (0.94-1.06) | 188177 | 1.00 (0.99-1.01) | 3644 | 1.00 (0.95-1.05) |
| Overweight (25.0-29.9 kg/m^2^) | 664 | 1.98 (1.72-2.28) | 67645 | 1.91 (1.89-1.94) | 1395 | 2.53 (2.31-2.77) |
| Obese (≥30.0 kg/m^2^) | 144 | 2.23 (1.58-3.15) | 22195 | 2.86 (2.79-2.93) | 362 | 4.24 (3.43-5.25) |
| **BMI categories (South Asian cut-offs)** |  |  |  |  |  |  |
| Underweight (<18.0 kg/m^2^) | 607 | 0.64 (0.57-0.71) | 33423 | 0.68 (0.67-0.69) | 543 | 0.61 (0.54-0.68) |
| Normal weight (18.0-23.0 kg/m^2^) | 1689 | 1.00 (0.93-1.07) | 146208 | 1.00 (0.99-1.01) | 2855 | 1.00 (0.95-1.05) |
| Overweight (23.0-27.0 kg/m^2^) | 1034 | 2.03 (1.82-2.25) | 96210 | 1.76 (1.74-1.78) | 1858 | 2.02 (1.89-2.17) |
| Obese (≥27.0 kg/m^2^) | 361 | 2.88 (2.32-3.57) | 46444 | 2.88 (2.83-2.93) | 870 | 3.91 (3.43-4.46) |
| **Trend (per 5 kg/m^2^)** | 3691 | 1.96 (1.81-2.11) | 322285 | 1.70 (1.69-1.71) | 6126 | 2.22 (2.10-2.34) |

^†^ Logistic regression models were adjusted for age, sex, area of residence, wealth index and highest educational attainment.
